# pykarambola: Minkowski tensor morphometry of 3D structures

**DOI:** 10.64898/2026.06.16.730752

**Authors:** Yajushi Khurana, Keisuke Ishihara

## Abstract

Three-dimensional biological morphologies encode functional and physiological state, yet the directional, orientational, and topological properties of these shapes are rarely captured by morphometric tools available for bioimage analysis. Minkowski tensors are mathematically rigorous tensor-valued measures that encode surface curvature and directionality for objects of arbitrary topology, with tensor eigensystems that directly quantify elongation axes and anisotropy. A C++ implementation, karambola (<u>1</u>), computes Minkowski tensors for triangulated surfaces but is inaccessible within Python-based bioimage workflows. Here we present pykarambola, a pip installable Python package that accepts NumPy arrays and standard mesh formats and returns Minkowski tensors, including derived anisotropy and orientation quantities. A high-level label-image API converts 3D integer arrays into per-object Minkowski tensors in a single call, making pykarambola directly compatible with the output of widely used segmentation tools. An optional Cython extension accelerates graph-traversal steps of mesh initialization for large-scale analyses. Benchmarked on 1,584 adrenal gland meshes, pykarambola reproduces all 121 C++ karambola output features to near-floating-point agreement and, in the pure-Python build, is 2.8× faster at 28^3^ and 1.5× faster at 64^3^ voxel resolution, with speedups primarily attributable to karambola’s sequential per-object file I/O. pykarambola is freely available as an open-source software package.

## Introduction

Modern fluorescence microscopy and volumetric segmentation routinely produce three-dimensional representations of biological structures (e.g., organelles, nuclei, cells, and tissue regions) at single-object resolution, enabling systematic morphometric analysis at scale. Quantifying the shapes of these structures requires descriptors that go beyond size: elongation, surface orientation, and topological complexity often carry direct biological meaning, reflecting organelle function, cell polarity, or tissue organization. Standard descriptors such as volume, surface area, sphericity, and inertia tensor eigenvalues (as computed in scikit-image (2)) capture the geometry of the mass distribution but not the directed properties of the surface itself: its local curvature, normal orientation, and their spatial distribution. Spherical harmonics-based descriptors encode richer structural information (3, 4), but are restricted to genus-zero surfaces and cannot describe objects with handles, tunnels, or enclosed voids, which arise in multi-component segmentations and topologically complex organelles. A shape descriptor that is simultaneously orientation-sensitive, topology-aware, and directly interpretable in terms of elongation and anisotropy is therefore needed.

Minkowski tensors provide this combination. The classical Minkowski functionals form a complete basis for continuous, additive, and motion-invariant scalar shape measures: by Hadwiger’s theorem, any such functional on convex bodies in ℝ^3^ is a linear combination of exactly four measures (5): volume *V*, surface area *A*, integrated mean curvature *H*, and Euler characteristic *χ*, a topological invariant equal to the number of connected components minus the number of handles plus the number of enclosed voids. These four scalars describe size, surface extent, bending, and topology, but carry no directional information: a sphere, a cigar-shaped ellipsoid, and a flat disk can share identical *V, A, H*, and *χ* while differing profoundly in elongation and orientation. Minkowski tensors extend this framework by accumulating directional contributions from surface geometry (6), yielding a family of tensor-valued shape measures 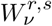 indexed by the geometric measure type *ν* (*ν* = 0: volume, *ν* = 1: surface area, *ν* = 2: mean curvature, *ν* = 3: Gaussian curvature weighted) and carrying total tensor rank *r* + *s* (6): scalars (*r* + *s* = 0), vectors (*r* + *s* = 1), rank-2 tensors (*r* + *s* = 2), and higher-rank tensors such as 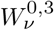 and 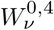. Minkowski tensors are physically interpretable. In addition, the eigenvalues and eigen-vectors of a tensor directly quantify the principal anisotropy magnitudes and orientation axes of the object. For example, the eigensystem of the rank-2 surface tensor 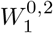, which accumulates outer products of surface normals weighted by area, directly quantifies elongation and anisotropy: the ratio of the smallest to the largest absolute eigenvalue defines the anisotropy index 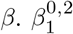 equals one for a perfect sphere and approaches zero for increasingly elongated or oblate shapes. A further layer of description comes from decomposing the Cartesian Minkowski tensors into irreducible representations under rotation, giving rise to the Minkowski structure metrics *q*_*ℓ*_ and *w*_*ℓ*_ (7, 8): rotationally invariant scalars that characterize *ℓ*-fold orientational order. In addition, Minkowski tensors are defined for triangulated surfaces of arbitrary topology (6), making them applicable without restriction to the full diversity of three-dimensional morphologies encountered in biological image analysis, including multi-component objects, structures with enclosed voids, and surfaces of non-trivial genus.

A C++ implementation, karambola (1), has been applied to characterize anisotropy in cellular foams, granular packings, and porous materials (6), but is not accessible within the Python bioimage ecosystem. Modern bioimage analysis pipelines are predominantly built around Python, with acquisition, segmentation, and quantification tools forming an interconnected ecosystem of libraries that operate on in-memory data structures. karambola accepts meshes exclusively in the karambola-native `.poly` format and the Object File Format (`.off`), and writes results to plain-text files; it is not distributed through a package manager and requires a C++ build toolchain to compile. Crucially, it provides no Python-compatible interface: since modern bioimage pipelines produce in-memory three-dimensional integer segmentation arrays rather than mesh files, integrating karambola requires serializing each segmented object to disk in a supported mesh format, executing the binary as an external process, and parsing the resulting plain-text output files back into Python, introducing fragile intermediates that are difficult to integrate and maintain.

Here we present pykarambola, a complete Python reimplementation of karambola that integrates Minkowski tensor morphometry directly into the Python bioimage ecosystem. pykarambola is installable via `pip install pykarambola`, accepts vertex and face arrays as NumPy arrays as well as mesh files in `.poly`, `.off`, `.obj`, `.glb`, and `.stl` formats, and returns results as plain Python dicts. It reproduces the full karambola feature set and additionally returns derived anisotropy indices and orientation quantities directly in its output, without the post-processing that karambola requires. A high-level label-image API (`minkowski_tensors_from_label_image`) accepts three-dimensional integer segmentation arrays directly, extracts a triangulated surface per object via scikit-image marching cubes (2), and returns per-object Minkowski tensors in a single call, making pykarambola immediately compatible with the integer label images produced by segmentation tools. An optional Cython extension accelerates graph-traversal steps of mesh initialization for large-scale analyses. pykarambola is freely available at https://github.com/Ishihara-SynthMorph/pykarambola and archived on Zenodo (https://doi.org/10.5281/zenodo.20418801).

## Methods

### Architecture of pykarambola

pykarambola is a complete Python reimplementation of the C++ karambola package (1), preserving the mathematical definitions and output conventions of karambola while replacing implementation-specific dependencies with Python-native equivalents. The port reproduces the logical structure of the C++ source: a `Triangulation` class holds the surface mesh and pre-computed per-triangle and per-edge geometry, a set of `calculate_w*` functions implements the Minkowski tensor formulae, and a command-line interface reproduces the behavior of the original binary. The C++ implementation depended on the GNU Scientific Library (GSL) for eigenvalue decomposition; pykarambola replaces this with `numpy.linalg.eigh`, so that all dependencies are installable as standard Python packages with no external C library required.

The package is structured around a pure-Python core that requires NumPy (9) and SciPy (10). SciPy is used in two places: connected-component labeling of mesh graphs (via `scipy.sparse.csgraph`) and evaluation of spherical harmonic basis functions for the Minkowski structure metrics (`msm`) (via `scipy.special`). Where pure-Python performance is insufficient for large meshes, an optional Cython extension accelerates the graph-traversal steps of mesh initialization; the package falls back to pure-Python implementations when the extension is not installed. To support direct processing of segmentation outputs, a high-level label-image API (`minkowski_tensors_from_label_image`) additionally requires scikit-image (2) for marching-cubes surface extraction; this dependency is intentionally optional so that the core mesh API remains lightweight, and it resolves automatically in standard bioimage Python environments where scikit-image is typically present.

pykarambola retains support for the two mesh formats accepted by C++ karambola: the karambola-native `.poly` plain-text polygon-mesh format and the Object File Format (`.off`). Three additional parsers for Wavefront OBJ (`.obj`), binary glTF (`.glb`), and STereoLithography (`.stl`) were added to broaden compatibility with mesh files produced by common 3D reconstruction and visualization tools.

The package ships with an extensive test suite (runnable via `pytest`) covering numerical agreement with C++ karambola across all output features, mesh quality validation for open, non-manifold, and degenerate surfaces, and correctness of the label-image and file-format APIs. Numerical correctness of the port was further validated by extensive benchmarking against the C++ karambola binary across a diverse set of test meshes, confirming near-floating-point agreement for all Minkowski scalars, vectors, and tensors.

### Acceleration strategy: NumPy vectorization and Cython

pykarambola achieves competitive runtime primarily through NumPy vectorization (9): all mesh data are stored as contiguous arrays and bulk operations replace Python-level loops over individual triangles or edges. During `Triangulation` construction, triangle normals, face areas, triangle centroids, edge lengths, and dihedral angles are precomputed via batched cross-products and array-wise norm computations on (*F* × 3) arrays, where *F* is the number of triangles and 3 reflects either the three Cartesian components (for normals and centroids) or the three edges per triangle (for edge lengths and dihedral angles). Each Minkowski tensor function then reduces these precomputed arrays to per-label scalar, vector, or matrix accumulators using `numpy.sum` over element-wise array products, with no Python-level loop over faces.

Two graph-traversal steps resisted vectorization: building the triangle-adjacency table (a hash map indexed by vertex-pair keys) and constructing the vertex-to-triangle lookup table. Both require data-dependent branching over irregular graph structures that cannot be expressed as bulk array operations. These two routines are therefore provided as an optional Cython extension (`_accel.pyx`) that uses typed memoryviews and eliminates interpreter overhead on the inner loops. When the extension is not installed, equivalent pure-Python implementations are used as a fallback.

### High-level Python API

pykarambola exposes two high-level functions that accept NumPy arrays or `Triangulation` objects and return plain Python dicts. The first, `minkowski_tensors`, accepts an (*V* × 3) vertex array (one row per vertex, three columns for *x, y, z* coordinates) and an (*F* × 3) integer face array (one row per triangle, three columns for vertex indices) and returns a dict whose string keys (`w000`, `w020`, `w020_eigvals, …`) map to NumPy scalars, vectors, or matrices. The `compute` argument controls which outputs are returned: `‘standard’` returns the 14 base tensors with their eigensystems; `‘all’` additionally returns the higher-order tensors `w103` and `w104` (including `w104` eigenvalues), the spherical Minkowski summary quantities `msm_ql` and `msm_wl`, and three derived scalars per rank-2 tensor. The derived scalars are `_beta` (anisotropy index, ratio of smallest to largest absolute eigenvalue), `_trace` (matrix trace), and `_trace_ratio` (trace normalized by the paired Minkowski scalar, e.g. 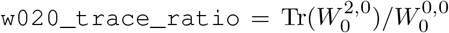; defined only for the 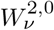 family). An explicit list of names may also be passed for selective computation. The boolean `compute_eigensystems` (default `True`) controls whether eigendecomposition is performed for rank-2 tensors; setting it to `False` omits the `*_eigvals` and `*_eigvecs` keys and avoids six `numpy.linalg.eigh` calls per label, which can reduce runtime for large batch jobs that do not require eigensystems. When the `labels` argument is provided as a per-face integer array, or set to `‘auto’` to detect connected components automatically via SciPy, the function returns a nested dict keyed by label. The optional `return_count=True` argument changes the return value to a `(results, n_objects)` tuple where `n_objects` is the number of connected components counted by vertex-adjacency graph traversal, useful for detecting multi-component objects within a single label. The `center` argument controls the reference point for position-dependent tensors (those with *r >* 0: the vector family 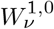 and the rank-2 family 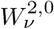 (6)): `None` uses the array origin, `‘reference_centroid’` reproduces the C++ karambola `-reference_centroid` flag, `‘centroid_mesh’` shifts each object to its volume-weighted center of mass before computing, and an explicit (3,) array applies a fixed shift; the boolean `center_per_label` (default `True`) determines whether centroid shifts are computed independently per labeled object or from the whole mesh. The second, `minkowski_tensors_from_label_image`, accepts a three-dimensional integer NumPy array in which each nonzero value identifies an object. For each label, it extracts a triangulated isosurface via scikit-image marching cubes and passes the resulting mesh to `minkowski_tensors`, returning a dict keyed by integer label with the same structure as the mesh API. The `spacing` argument propagates physical voxel dimensions to all length, area, and volume quantities, ensuring results are reported in physical units and are comparable across datasets acquired at different resolutions. The `autolabel=True` flag treats the array as a binary mask and delegates connected-component splitting to the mesh API.

## Results

### Runtime benchmark

We benchmarked pykarambola against the reference C++ karambola binary on two mesh sets. First, synthetic icospheres at subdivision levels 5–10 (10,242 to 10,485,762 vertices) were generated in pure NumPy by iterative triangular subdivision of a regular icosahedron. Both the pure-Python and Cython-accelerated builds were timed end-to-end (file parse plus compute) on these meshes. Second, the AdrenalMNIST3D dataset, a standardised biomedical benchmark (11) comprising 1,584 adrenal gland meshes at two voxel resolutions (28^3^ and 64^3^), was used to measure performance on realistic biological meshes. All measurements were performed on a single core of an Apple M4 processor (macOS 15.7.3, Python 3.13, pykarambola v0.5.0).

For the icospheres, the Cython-accelerated build (`pykarambola[accel]`) was generally faster than `karambola` (1.0–1.15× speedup for mesh sizes up to 2.6M vertices; Supplementary Figure S1b), with both within 3% of each other at the largest mesh size tested (10.5M vertices). The pure-Python build ran at 0.79–0.97× of `karambola`’s speed across all mesh sizes (Supplementary Figure S1b). Within `pykarambola`, Cython acceleration reduced compute time by 1.2–1.3× relative to the pure-Python build, with the benefit most pronounced at larger mesh sizes.

On AdrenalMNIST3D, `pykarambola` was 2.8× faster than C++ `karambola` at 28^3^ resolution and 1.5 ×faster at 64^3^ resolution across all 1,584 meshes. The C++ `karambola` timing includes sequential file I/O (one result folder per mesh written to disk), whereas the `pykarambola` timing covers only in-memory computation; the file I/O over-head of the C++ binary accounts for most of the speedup observed on the adrenal dataset.

### Numerical accuracy

To assess numerical accuracy, we compared all 121 output features of `pykarambola` against C++ `karambola` on the full AdrenalMNIST3D dataset (11) (1,584 meshes at both 28^3^ and 64^3^ resolutions). The 121 quantities comprise: 4 Minkowski scalars, 12 vector components, 36 rank-2 tensor upper-triangle elements (each rank-2 tensor is a symmetric 3 × 3 matrix with 6 independent entries), 18 rank-2 tensor eigenvalues, 27 components of the rank-3 tensor 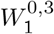 (`w103`), 6 eigenvalues of 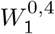 (`w104`), and 18 spherical Minkowski summary values (`msm_ql`, `msm_wl`).

Tensor eigenvectors are computed by `pykarambola` but are excluded from direct comparison: eigenvectors are defined only up to sign and, in the presence of degenerate eigenvalues, up to arbitrary rotation within the degenerate subspace; the eigenvalues fully characterize the spectral geometry of each tensor.

Across all 121 features, all 1,584 meshes, and both resolutions, `pykarambola` and `karambola` outputs fall on the identity line in each scatter plot (Supplementary Figure S2). Residual discrepancies are consistent with floating-point summation order differences between the C++ and Python implementations and with the substitution of the GNU Scientific Library eigenvalue solver with `numpy.linalg.eigh`. The maximum mean absolute difference was 1.6 × 10^−4^, observed for 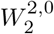 diagonal components at 64^3^ resolution. Scalar, vector, rank-3, and rank-4 features agree to better than 5 × 10^−9^ on average; the larger discrepancies in the rank-2 tensor family are consistent with summation-order sensitivity arising from their coordinate-product terms, whose magnitude scales with mesh size.

The spherical Minkowski summary quantities (`msm_ql` and `msm_wl`, *l* = 0, …, 8) are rotationally invariant scalars derived from the surface normal distribution: *q*_*l*_ measures *l*-fold orientational order and *w*_*l*_ captures its phase structure. They show a distinct accuracy pattern (Supplementary Figure S2). For `msm_ql` and even-*l* `msm_wl`, both implementations agree to floating-point precision. For odd-*l* `msm_wl` (*l* = 1, 3, 5, 7), a Wigner 3*j* parity selection rule (7, 12), a symmetry constraint on angular momentum coupling coefficients, requires *w*_*l*_ = 0 exactly. `pykarambola`’s implementation uses the Racah formula (12), a closed-form algebraic expression for Wigner 3*j* symbols that evaluates them exactly using integer arithmetic, and applies the parity rule analytically, returning exactly zero. C++ `karambola` instead uses `gsl_sf_coupling_3j`, which evaluates numerically without an explicit parity check, producing spurious residuals of order 10^−8^–10^−7^ on the AdrenalMNIST3D meshes.

### Label-image API walkthrough

The label-image API (`minkowski_tensors_from_label_image`) accepts integer segmentation arrays directly, converting each labeled region to a triangulated mesh and returning Minkowski tensors in a single call (Figure 1). To demonstrate the API, we applied it to a publicly available 3D fluorescence segmentation of a human induced pluripotent stem cell (hiPSC) in early anaphase/telophase from the Allen Cell Collection (4). The segmentation provides a two-channel binary mask at isotropic voxel spacing of 0.108 *µ*m: one channel for the DNA and one for the cell membrane.

**Fig. 1.**
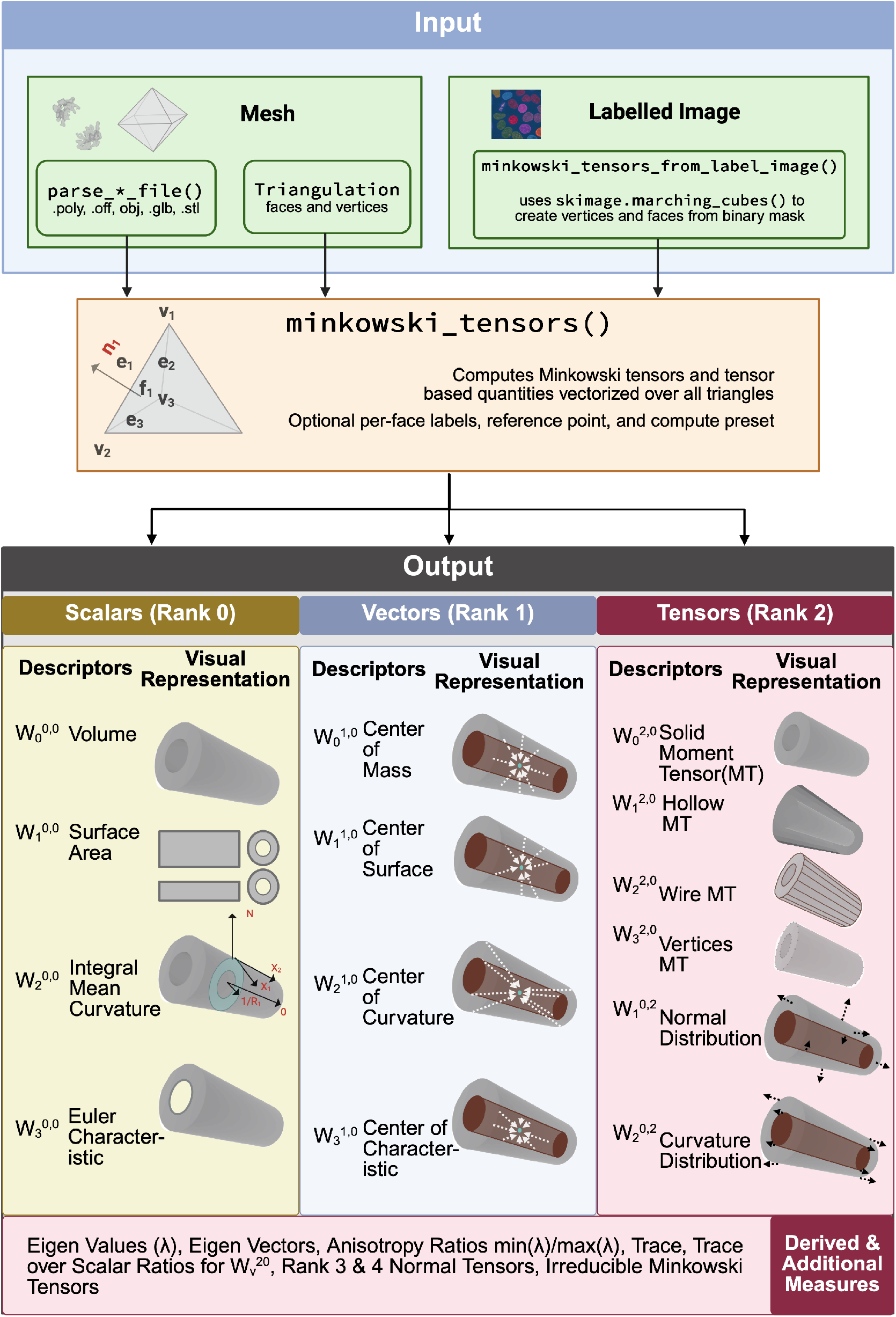
Computation pipeline of pykarambola. Three entry paths converge at the shared core (`minkowski_tensors`): mesh files (`.poly`, `.off`, `.obj`, `.glb`, `.stl`) via format-specific parsers (`parse_*_file`); `Triangulation` objects (vertex and face arrays) passed directly; and 3D integer label images via `minkowski_tensors_from_label_image`, which extracts a triangulated isosurface per label using `skimage.marching_cubes`. The core precomputes per-triangle geometry vectorized over all triangles and evaluates the Minkowski tensors (6). Outputs are organized into three families (physical interpretations after appropriate normalization; see (6) for exact expressions): **Scalars** 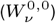: volume, surface area, integrated mean curvature, and Euler characteristic; **Vectors** 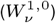: curvature-weighted centroids; and **Rank-2 tensors** (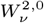: solid, hollow, wire, and vertex moment tensors; 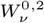 normal and curvature distribution tensors). With `compute=‘standard’` (default), eigenvalues and eigenvectors are returned for each rank-2 tensor. With `compute=‘all’`, additionally returned are: anisotropy index *β* = *λ*_min_*/λ*_max_, trace, and trace-over-scalar ratio per rank-2 tensor; rank-3 and rank-4 normal tensors; and irreducible Minkowski structure metrics.

Supplementary Figure S4 illustrates how the choice of labeling scheme directly determines what the tensors measure, using the DNA channel as an example. In case (i), the binary DNA mask is cast to an integer array (`dna_mask.astype(int)`) and passed with the default `autolabel=False`; because all non-zero voxels share label value 1, the function returns one tensor set describing the aggregate geometry of all chromosomal material (*V* = 110 *µ*m^3^, *A* = 435 *µ*m^2^, *χ* = 2 (a topologically closed, simply connected surface), 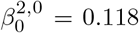). In case (ii), `scipy.ndimage.label(dna_mask)` first assigns a unique integer to each connected component; the resulting three-valued label image is passed with `autolabel=False` (default), and the function returns three independent tensor sets, one per DNA object. Equivalently, case (ii) can be achieved in a single call by passing the binary mask with `autolabel=True`, which instructs the function to detect connected components on the mesh internally without any preprocessing; the per-object tensors are numerically identical, as confirmed by matching per-object volumes and *β* values. Case (ii) answers a fundamentally different morphometric question than case (i): object 1 (*V* = 56 *µ*m^3^,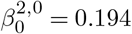) is approximately twice the volume of objects 2 and 3 (*V* ≈ 27 *µ*m^3^ each), which are more elongated (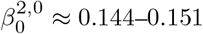). Objects 2 and 3 occupy non-overlapping axial ranges (Z slices 40–72 and 78–110) and together form segments of the same daughter chromosomal mass, consistent with a cell in early anaphase/telophase. The DNA objects are substantially more anisotropic than the cell membrane (DNA 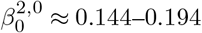 versus cell membrane 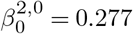 lower *β* indicates greater shape anisotropy).

This example demonstrates a general principle: identical binary image data yields fundamentally different morphometric results depending on label assignment, and the correct choice depends on the biological question. Whole-structure morphometry (case i) is appropriate when the aggregate shape is the object of interest; per-object morphometry (case ii) is required when individual instances must be resolved, as in cell counting, organelle tracking, or shape clustering. The full worked example including mesh visualisation and Minkowski tensor additivity is provided in `multilabel_image_workflow.ipynb`, available in the project GitHub repository.

## Discussion

pykarambola makes Minkowski tensor morphometry directly accessible within the Python bioimage ecosystem by replacing a C++ binary with a pip-installable package that accepts NumPy arrays and returns plain Python dicts. Beyond making the tool pip-installable, the reimplementation extends input support to Wavefront OBJ, binary glTF, and STL files and adds a high-level label-image API that converts 3D segmentation outputs into per-object tensors in a single call. The hiPSC anaphase example in the Results section demonstrates that this analysis is accessible directly from the integer label arrays produced by standard segmentation tools, without any intermediate mesh preparation. The benchmark results show that on icospheres, the Cython-accelerated build was generally faster than C++ karambola (1.0–1.15× speedup for mesh sizes up to 2.6M vertices, with both within 3% at the largest mesh tested), while the pure-Python build ran at 0.79– 0.97× of karambola’s speed across all mesh sizes. On the AdrenalMNIST3D dataset, pykarambola was faster than C++ karambola (2.8× at 28^3^, 1.5× at 64^3^), primarily because C++ karambola writes one result folder per object to disk. The NumPy vectorization of the Minkowski tensors over all triangles in a single array operation avoids per-triangle Python overhead, keeping the compute step comparable in speed to the C++ implementation. The optional Cython extension targets only the graph-traversal steps of mesh initialization, which resist vectorization; its benefit is therefore most pronounced for large meshes and batch analyses where initialization cost dominates. For interactive or small-scale use, the pure-Python build is sufficient and requires no compiled extension. A parallel-processing workflow for large-scale batch analyses using a lightweight Python library for transparent parallelization across CPU cores is also available and demonstrated in the project repository, saving additional computation time for large imaging datasets.

The primary limitation of pykarambola, inherited from karambola, is that it operates exclusively on triangulated surface meshes. For label-image inputs, the surface is extracted by marching cubes, which introduces a discretization artifact: at typical fluorescence microscopy voxel sizes, the mesh is a piecewise-planar approximation of the true surface. For smooth objects, tensor values converge asymptotically as voxel resolution increases, but rank-2 surface tensors are subject to an orientation bias from the anisotropy of the digital grid that can be significant at typical fluorescence microscopy resolutions (13); an alternative Voronoi-based approach that is asymptotically unbiased for binary voxel inputs without requiring surface reconstruction has recently been proposed (14). A further limitation applies to the direct mesh API, where users may encounter open or non-manifold meshes in their biological datasets. In such cases pykarambola issues a warning, noting that for open surfaces, volume-dependent quantities (`w000`, `w020`) are set to NaN, while non-manifold geometries render all results unreliable; users should therefore verify mesh quality before interpreting tensor outputs. The label-image API is not affected, as marching cubes always produces closed, manifold surfaces by construction.

Future development directions include support for additional Minkowski tensor families beyond those available in karambola and a streaming batch-processing mode for datasets too large to hold in memory simultaneously. pykarambola’s speed makes it well-suited for real-time feedback as users explore and annotate 3D segmentations. Integration into interactive analysis environments is a particularly attractive direction, for example as a napari (15) plugin that visualizes tensor outputs alongside raw image data and allows users to explore morphometric results in real time. Systematic descriptors of three-dimensional shape have been absent from the bioimage ecosystem. By closing that gap, pykarambola brings Minkowski tensor morphometry to the scale and diversity of modern biological imaging datasets, enabling studies of organelle geometry, cell shape, and tissue organization that were previously impractical.

## Software availability

pykarambola is released under the BSD 3-Clause License and is freely available at https://github.com/Ishihara-SynthMorph/pykarambola. The package can be installed via the Python Package Index (PyPI) with `pip install pykarambola`. Version 0.5.0, the version described in this paper, is archived on Zenodo at https://doi.org/10.5281/zenodo.20418801. Core dependencies are NumPy and SciPy; optional dependencies include Cython (for acceleration) and scikit-image (for label-image API).

## Data availability

All datasets used in this study are publicly available. The AdrenalMNIST3D data is part of MedMNIST (11) and can be downloaded via the `medmnist` Python package. The Allen Cell Collection segmentations are publicly available through the Allen Institute for Cell Science (4).

## ACKNOWLEDGEMENTS

pykarambola was developed by directly referencing the source code of karambola, the C++ reference implementation by Schaller, Kapfer, and Schröder-Turk (1). The authors of karambola kindly agreed to pykarambola being released under the BSD 3-Clause License. This work was supported in part by an American Heart Association Career Development Award (grant 25CDA1432232) to K.I.

## AI tools disclosure

Claude Opus 4.5 and Claude Sonnet 4.6 (Anthropic) were used during this project for software and code development (Python port, refactoring, and test scaffolding) and for manuscript drafting and editing. AI-assisted outputs were reviewed, edited, and validated by the human authors before inclusion.

## AUTHOR CONTRIBUTIONS

Y.K. and K.I. contributed equally to Software. Y.K.: Methodology, Validation, Visualization, Writing – original draft. K.I.: Conceptualization, Funding acquisition, Project administration, Supervision, Writing – review & editing.

## Supplementary Material

**Fig. S1.**
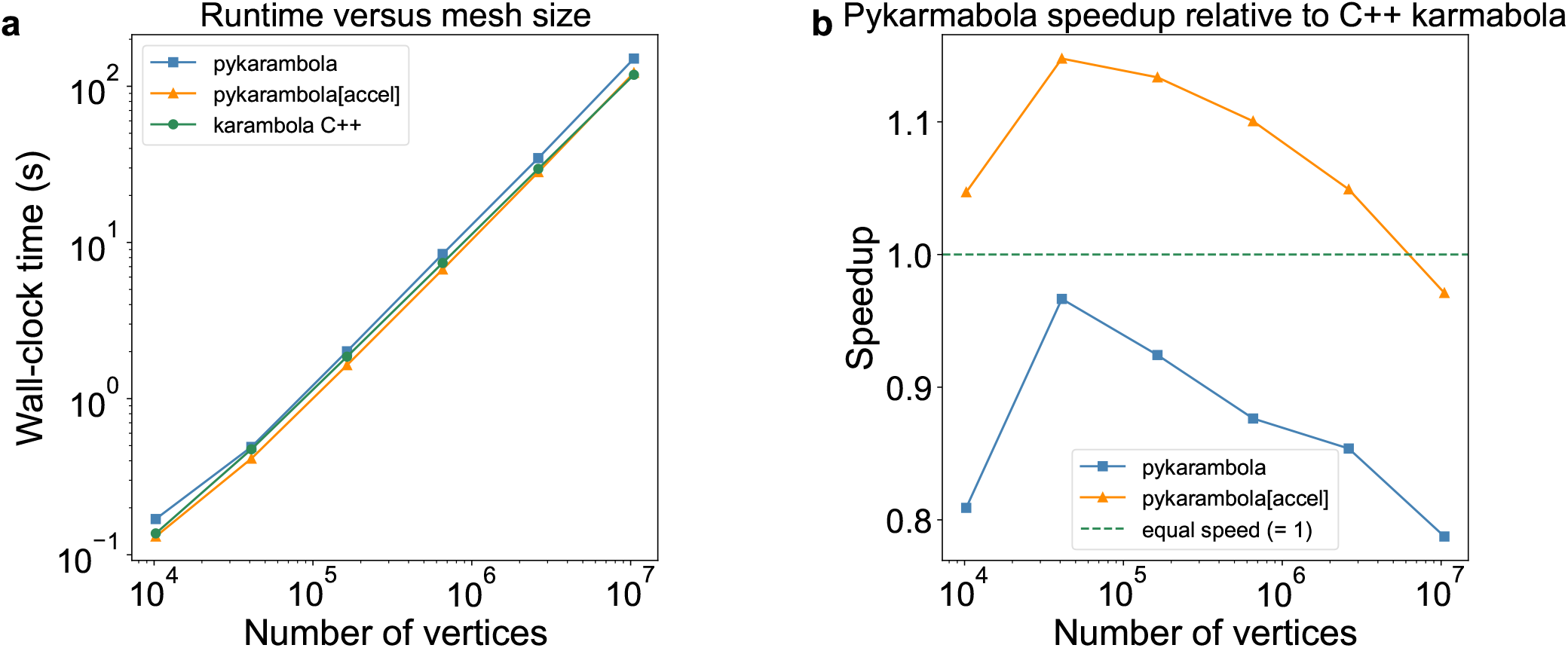
Runtime benchmark on synthetic icospheres (subdivision levels 5–10; 10,242–10,485,762 vertices). **Left**: absolute wall-clock time (log-log scale) for end-to-end execution (mesh parse + Minkowski tensor compute) for C++ karambola (green), pykarambola pure-Python (blue), and pykarambola Cython-accelerated (orange). **Right**: speedup ratio (karambola time / pykarambola time); values above the dashed line (= 1) indicate pykarambola is faster.

**Fig. S2.**
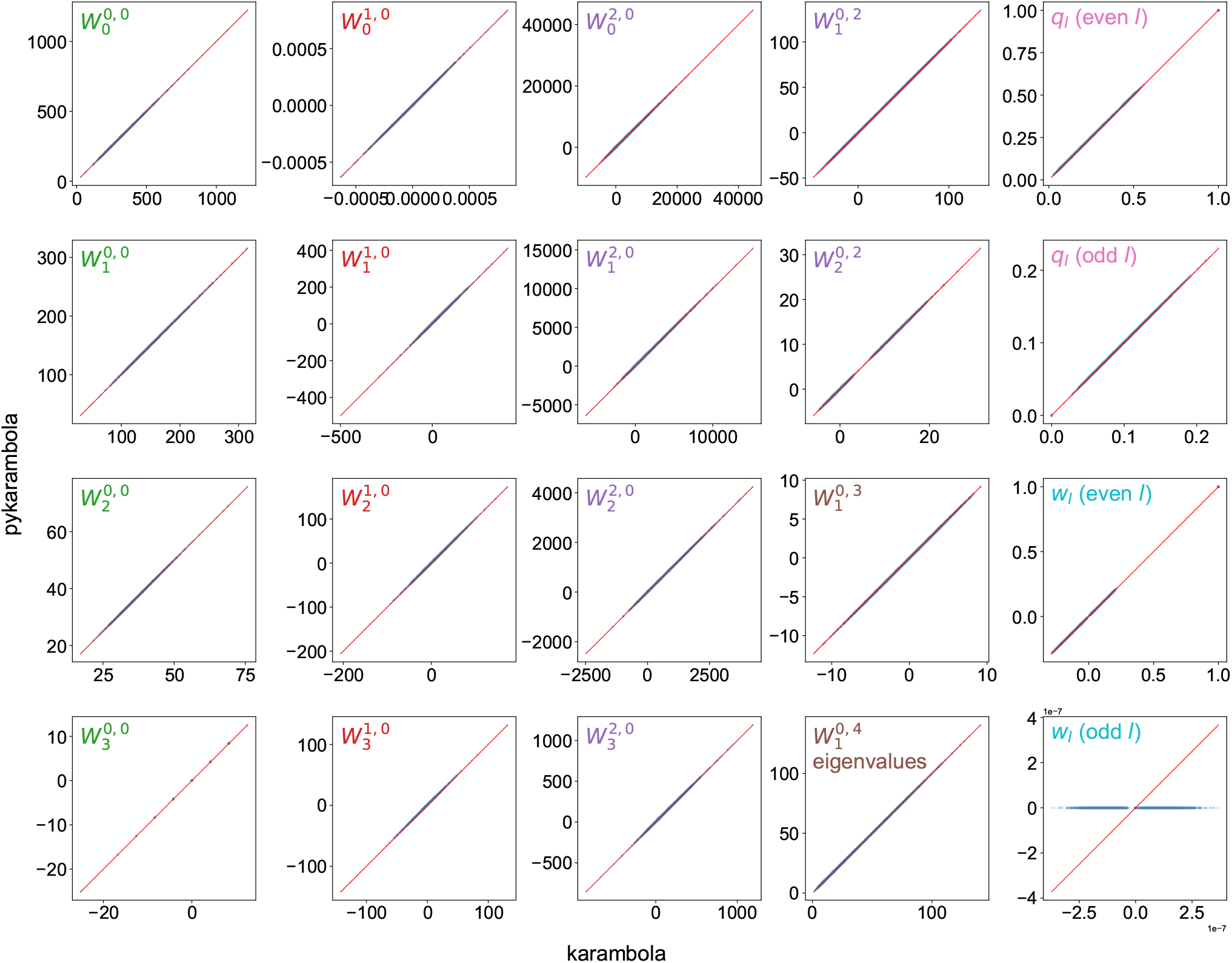
Numerical accuracy of `pykarambola` relative to C++ `karambola`. Each panel shows a scatter plot of values from identical analyses run in C++ (`karambola`, *x*-axis) and Python (`pykarambola`, *y*-axis) on the AdrenalMNIST3D dataset (1,584 meshes, 28^3^ voxel resolution). Columns correspond to feature groups: scalars 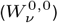, vectors 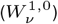, rank-2 tensors 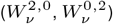, and higher-rank quantities (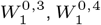 eigenvalues, and spherical Minkowski summary quantities *q*_*l*_ and *w*_*l*_). Panel labels are shown in the top-left corner of each panel, colored by feature group. The red diagonal indicates perfect agreement; all points fall on this line except odd-*l* `msm_wl`, which `pykarambola` returns as exactly zero while `karambola` produces small spurious residuals (see text). Equivalent agreement is observed at 64^3^ resolution (Supplementary Figure S3).

**Fig. S3.**
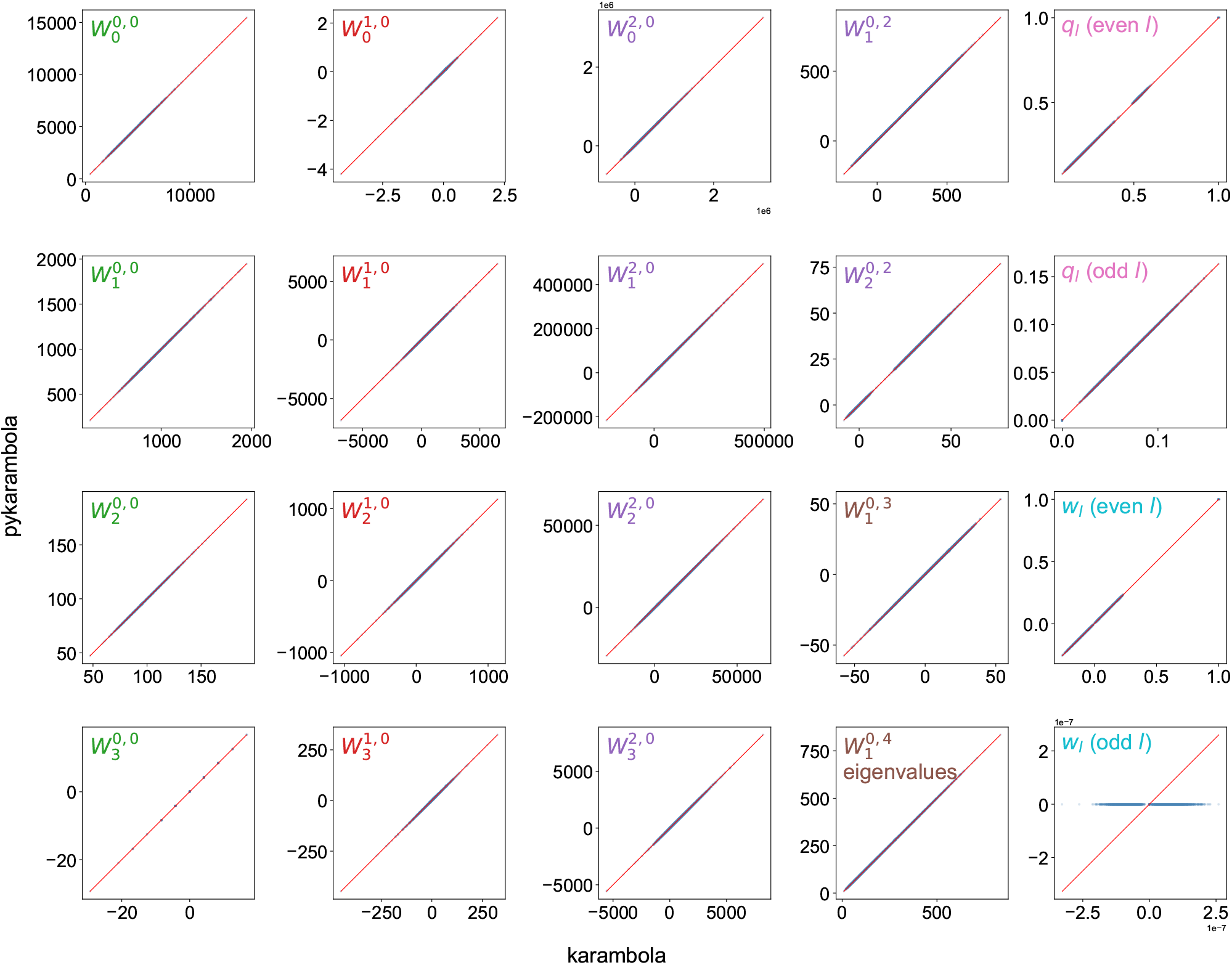
Numerical accuracy of `pykarambola` relative to C++ `karambola` at 64^3^ voxel resolution. All panels and groupings are identical to Supplementary Figure S2.

**Fig. S4.**
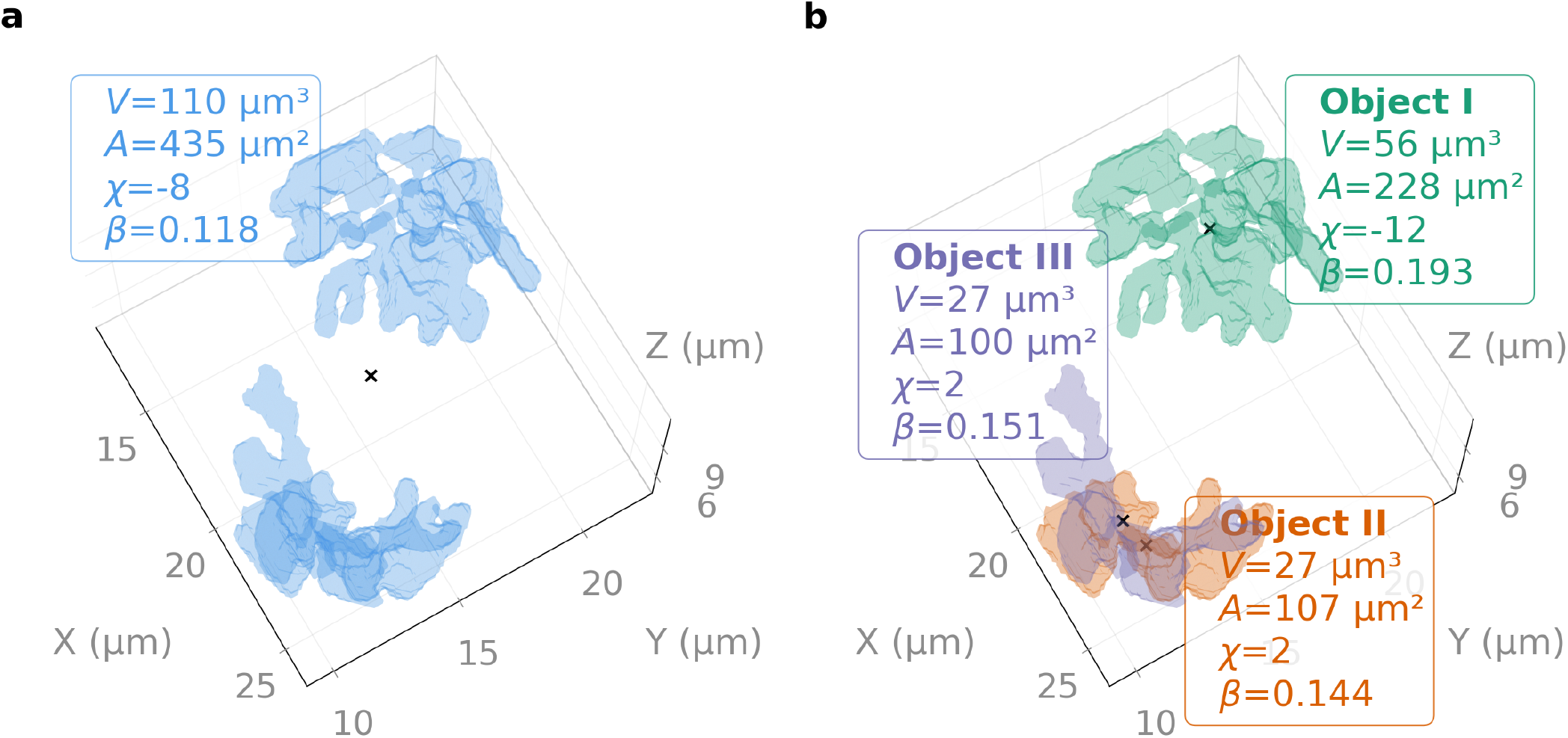
Label-image API applied to the DNA channel of a 3D fluorescence microscopy segmentation of a hiPSC in anaphase/telophase (Allen Cell Collection (4); isotropic voxel size 0.108 µm). Per-object Minkowski tensors report volume (*V*), surface area (*A*), Euler characteristic (*χ*), and anisotropy index (*β*), with object centroids marked (×). (**a**) The entire DNA channel assigned as a single label; one tensor set describes the aggregate of all three DNA objects. (**b**) The binary DNA mask passed with `autolabel=True`; pykarambola detects three connected components internally and returns three independent tensor sets, one per DNA object. Objects 2 and 3 occupy non-overlapping axial ranges (Z slices 40–72 and 78–110, respectively) and are likely segments of the same daughter chromosomal mass.

